# Identification of Aggregation Pheromone in *Odontothrips loti* (Thysanoptera: Thripidae)

**DOI:** 10.1101/2024.03.25.586550

**Authors:** Liu Yanqi, Luo Ying Ning, Liu Chang, Ban Liping

## Abstract

Pheromone trapping is an interspecific, active at low concentrations, eco-friendly pest management strategy that has been widely used for population monitoring. Pheromones have only been validated for a few species in Thysanoptera and the pheromone components of *Odontothrips loti* Haliday are still unclear. In our study, we have identified (R)-lavandulyl (R)-2-methylbutanoate from male *O. loti* headspace volatiles by gas chromatography-mass spectrometry (GC-MS), which was structurally similar to the known reported aggregation pheromone of *Megalurothrips sjostedti*. Y-tube olfactometer assays showed that both male and female adults *O. loti* were significantly attracted by synthetic (R)-lavandulyl (R)-2-methylbutyrate, implicating as an aggregation pheromone. Additionally, electroantennogram responses of *O. loti* increased with increasing doses of synthetic (R)-lavandulyl (R)-2-methylbutyrate. This is the first report of a male-produced aggregation pheromone in *O. loti*, from genus *Odontothrips*. The discovery of aggregation pheromone of *O. loti* as a primary pest of alfalfa will provide the possibility of monitoring and early warning in alfalfa grass fields and is expected to be used for integrated management for *O. loti*.

## Introduction

*Odontothrips loti* Haliday (Thysanoptera: Thripidae) is an oligophagous and devastating pest that threatens alfalfa (*Medicago sativa*) due to their incredible fecundity, insidiousness, and potential as vectors of destructive viruses, such as alfalfa mosaic virus, affecting the yield and quality of alfalfa hay [1]. Thrips are notoriously difficult to manage, in part because of serious overlapping generations, high outbreak frequency, and rapidly developing resistance due to overuse of chemical pesticides [2,3]. The *O. loti* has developed resistance to insecticides such as neonicotinoids, pyrethroids, and benzamides [4]. Although the application of broad-spectrum insecticides temporarily controls thrips abundance, it fails to mitigate the spread of viruses carried by thrips in the field [5]. Moreover, adverse effects on natural predators result in population resurgences that complicate thrips control [5]. Consequently, current research is primarily focused on developing effective and environmentally friendly strategies for thrips control.

Behavioral control of herbivores is a promising alternative and has been widely used to monitoring population dynamics of pests [6]. It is utilized to set traps using the visual and/or olfactory cues associated with their foraging and reproductive behaviors. Studies have shown that western flower thrips (*Frankliniella occidentalis*) prefers yellow and blue as visual signals in the laboratory and in the field [6,7]. Nonetheless, visual cues were often susceptible to distance limitations, and the long-distance localization relied more on olfactory cues, i.e., semiochemicals [7].

Some semiochemicals, including allelochemicals and pheromones, have demonstrated efficacy in orienting the behavior of insects both in natural environments and controlled settings like greenhouses. They serve as an integral component of numerous integrated pest management (IPM) programs. Allelochemicals play a crucial role in facilitating communication between different species, while pheromones are involved in communication within the same species. It has been shown that feeding triggers the production of herbivore-induced plant volatiles (HIPVs) [8,9], which vary with plant development and environmental conditions [10]. These compounds function to repel herbivores and pathogens or attract pollinators and seed dispersers to enhance reproduction [11,12], or even priming defense responses in the neighboring unattacked plants [13]. Scientists have named semiochemicals according to the responses they elicit from insects, such as attractants, repellents, arrestants [14]. Of these, attractants have been most maturely used in the “Push-Pull” strategy. For example, the combined utilization of attractants and entomopathogenic fungi has resulted in great savings for both labor and biopesticide dosage costs [15]. The combination of aggregation pheromone and alarm pheromone is successful in thrips control in greenhouses [16]. Therefore, research on the interactions between semiochemicals and pests is essential in developing effective methods to control thrips populations.

As early as the 1980s, researchers were drawn to the phenomenon of regular aggregations of male thrips, wherein females typically join these gatherings for mating purposes before departing [17,18]. Such male aggregations are commonly observed in plant-feeding populations of Thripidae and have been scientifically proven to be facilitated by aggregation pheromones emitted by the males. [19-24]. At present, thrips aggregation pheromones that have been studied in *F. occidentalis* [19], *F. intonsa* [20], *Megalurothrips sjostedti* [23], *M. usitatus* [24], *Thrips palmi* [21], *Dendrothrips minowai* [25]. The components are mostly terpene esters or monoterpene alcohols containing chiral carbon atoms. Exploring male-released aggregation pheromones could increase our understanding of the complexity of intra-population communication and be used in thrips management. For example, the aggregation pheromone of *F. occidentalis* was used to develop commercial lures [26]. In-field application of aggregation pheromones generally increases trapping up to 1-2 times, and even up to 7 times [6,24,25].

Currently, *O. loti* has emerged as a prominent pest species in alfalfa grass fields, garnering increasing attention. The aggregated distribution on alfalfa has been previously documented [27]. However, the active pheromones produced by *O. loti* have yet to be reported. The objectives of our study were to identify unknown pheromone components of *O. loti* and elucidate their roles, aiming to provide a valuable tool for the detection and management of commercial and quarantine pests.

## Result

### 2.1 Identification of headspace volatiles released by *Odontothrips loti*

The active volatile compounds from adult and second instar nymph were extracted using the solid-phase microextraction (SPME) method, followed by separation and identification through gas chromatography-mass spectrometry (GC-MS). A detailed comparison of TIC chromatograms revealed a male-specific compound (peak a) present at 27.37 min and a larvae-specific compound (peak b) present at 26.70 min (Figure 1, 2). Mass spectra analysis using the NIST mass spectral library identified the male-specific compound as (R)-lavandulyl (R)-2-methylbutanoate and the larvae-specific compound as (-)-Germacrene D. We further validated and characterized both compounds by comparing their retention times and mass spectra with authentic standards analyzed under the same conditions. Both native *O. loti* components had identical retention times to the standards, and the mass spectra were superimposable (Figure 1, 2).

**Figure 1.**
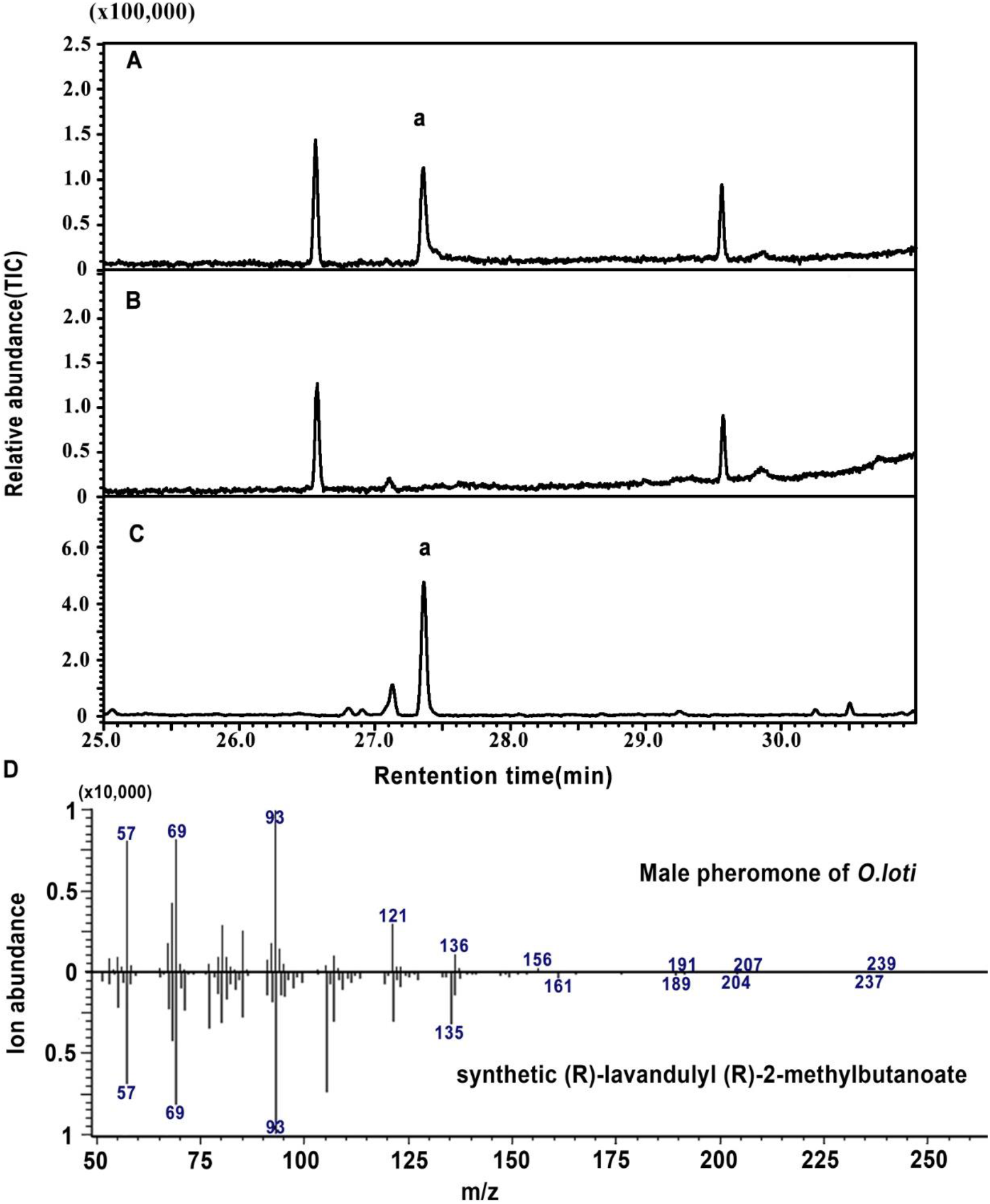
Representative chromatograms of the headspace volatiles collected from *Odontothrips loti* adult males (A), adult females (B), as well as synthetic (R)-lavandulyl (R)-2-methylbutanoate on a HP-5MS column. The labeled compound is (R)-lavandulyl (R)-2-methylbutanoate (a) Each experiment was performed in at least three biological replicates, with headspace volatiles collected from 10 insects per replicate, and all sample insects were used for only one collection. Electrospray ionization mass spectra of male-specific pheromone and their synthetic components.

**Figure 2.**
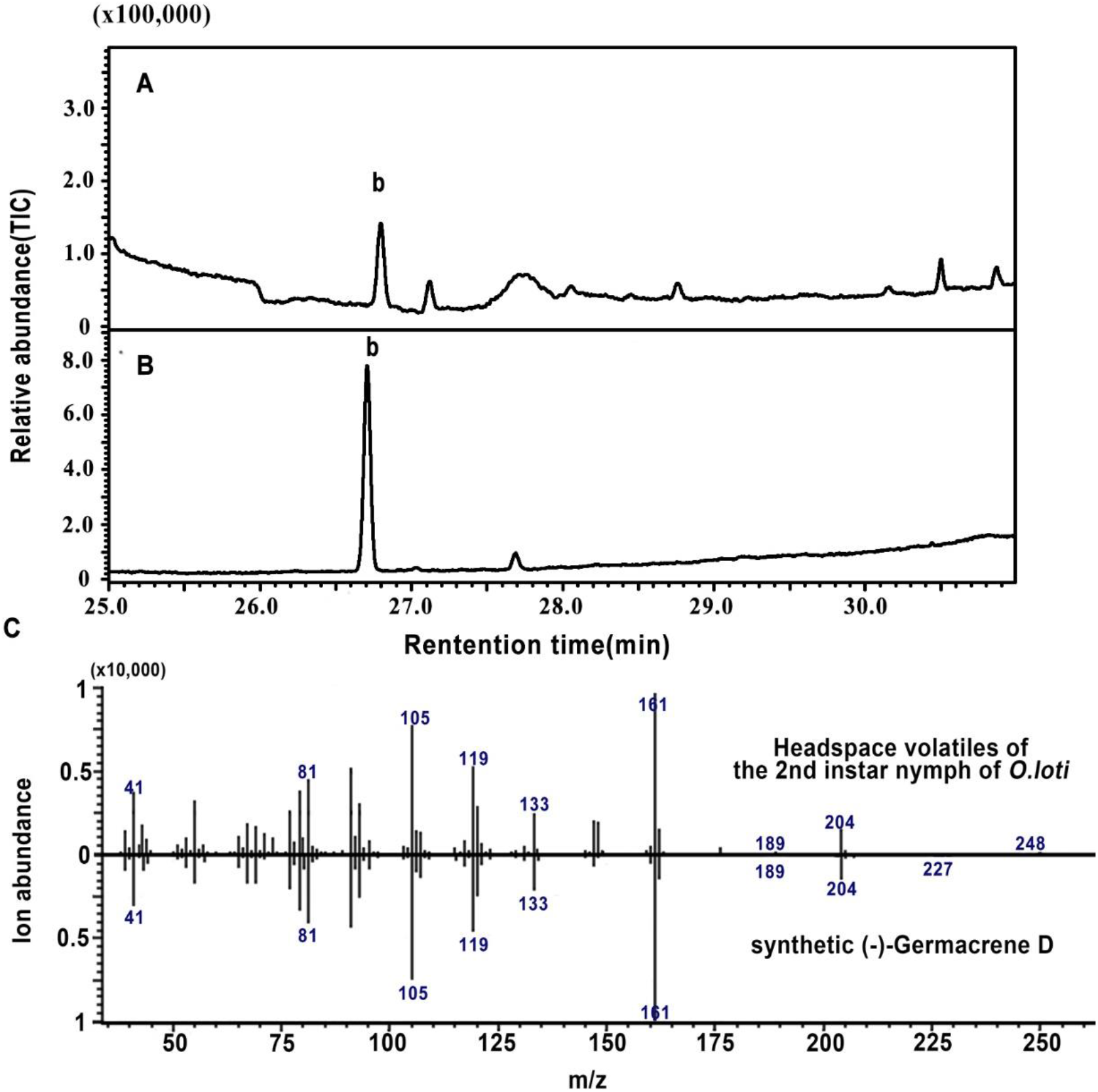
Representative chromatograms of the headspace volatiles collected from *Odontothrips loti* 2nd instar nymphs (A) and synthetic (-)-Germacrene D (B) on a HP-5MS column. The labeled compound is (-)-Germacrene D (b). Each experiment was performed in at least three biological replicates, with headspace volatiles collected from 10 insects per replicate, and all sample insects were used for only one collection. (C) Electrospray ionization mass spectra of 2nd instar-specific component and their synthetic components.

### 2.2 Behavioral response of *Odontothrips loti* to synthetic compounds

Due to the presence of two chiral carbon, there are four enantiomers for (R)-lavandulyl (R)-2-methylbutanoate. Y-tube olfactometer assays showed that male *O. loti* was significantly attracted to synthetic (R)-lavandulyl (R)-2-methylbutanoate but not to other enantiomers (Figure 3A). Further tests proved that both male and female adults *O. loti* demonstrated strong attraction towards synthetic (R)-lavandulyl (R)-2-methylbutyrate, as the test concentration increased from 0.5 µg/µl to 2 µg/µl. It is apparent that male *O. loti* responded at lower concentrations (0.5 µg/µl) with greater sensitivity than the female adults (Figure 3B). However, no differences were found between virgin females and females.

**Figure 3.**
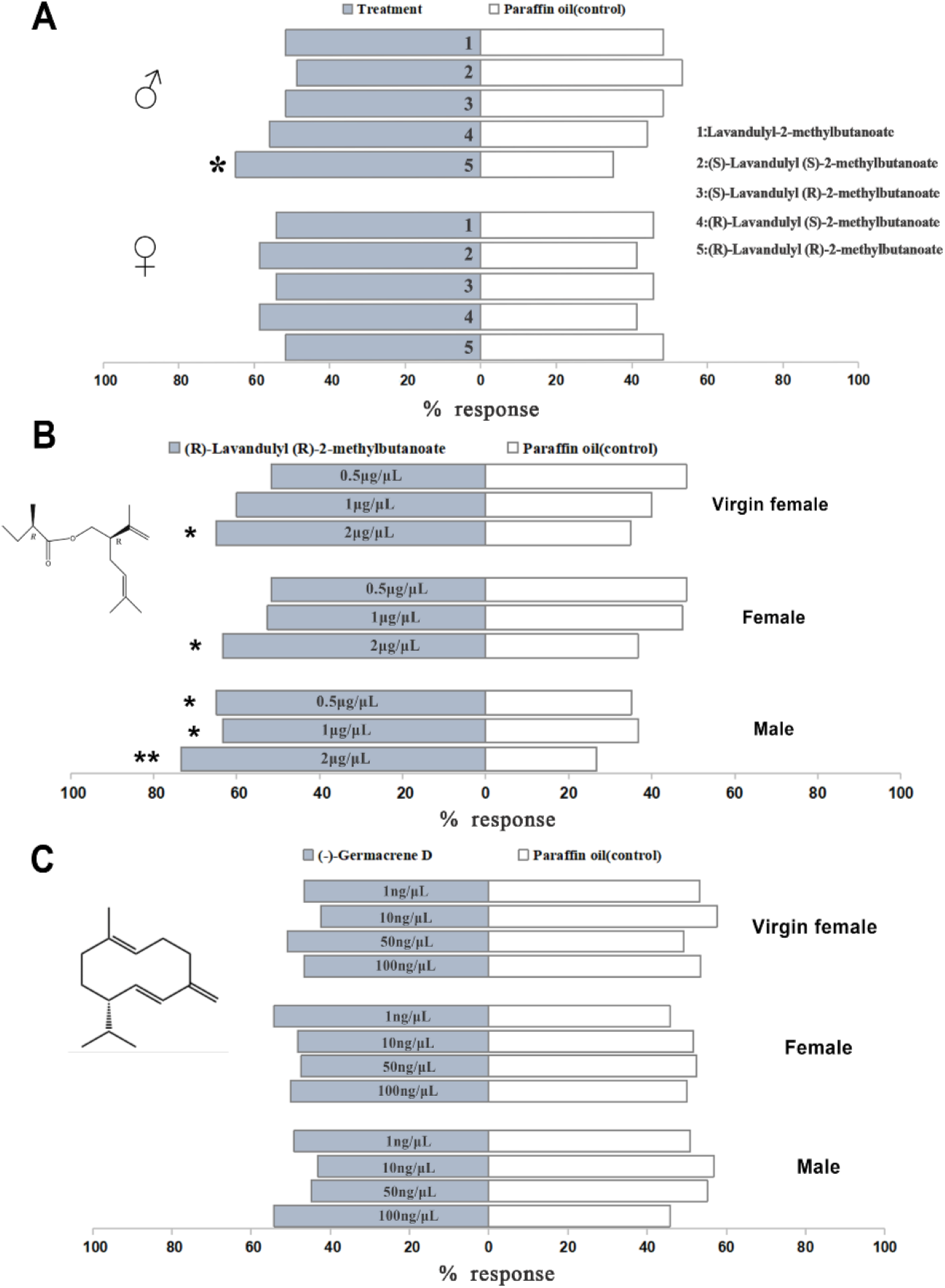
(A) Behavioral response of adult *O. loti* to synthetic lavandulyl-2-methylbutanoate and four enantiomers at 0.5μg/μl. (B) Behavioral response of adult *O. loti* to synthetic (R)-lavandulyl (R)-2-methylbutanoate at three doses. (C) Behavioral response of adult *O. loti* to synthetic (-)-Germacrene D at four doses. Sixty individuals per sex were tested in each treatment. Asterisks indicate a statistically significant difference by chi-squared test (**P < 0.01; *P < 0.05).

**Figure 4.**
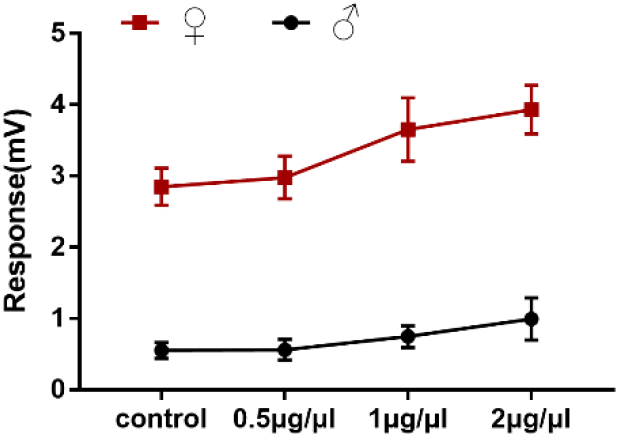
Electroantennogram responses of adult *O. loti* to synthetic (R)-lavandulyl (R)-2-methylbutanoate at three doses (n=10). Values are mean ± SE.

Regarding (-)-Germacrene D, the compound identified in the second instar nymph, when setting up four concentration gradients ranging from 1ng/µl to 100ng/µl of (-)-Germacrene D, neither male nor female adults *O. loti* exhibited any noticeable responses (Figure 3C).

### 2.3 Electrophysiological response of adult *O. loti* to synthetic (R)-lavandulyl (R)-2-methylbutanoate

Based on the strong behavioral response of *O. loti* to synthetic (R)-lavandulyl (R)-2-methylbutanoate, we conducted electrophysiological studies on adult male and female *O. loti* individuals to this compound. The EAG response of adult males and females to synthetic (R)-lavandulyl (R)-2-methylbutanoate generally increased as doses increased, and no significant difference within the designated concentration range.

## Materials and methods

### 3.1 Insect rearing

The adult *O. loti* used in this study were collected from *M. sativa* field grown in China Agricultural University, Beijing, China (40°1′42″ N, 116°16′43″ E) in July 2021. Thrips were reared in the laboratory with alfalfa leaves at 26±1°C, 60±5% RH under a photoperiod of 16 h light/8 h dark. During the third instar stage, thrips were individually selected and reared separately in 10ml centrifuge tubes, the bottoms of which were covered with dry toilet paper and lentil pieces. Virgin females come from groups that are reared separately at 3-5 days of age. Other adults from 3-5 days old rearing mixed groups [21].

### 3.2 Collection of headspace volatiles from *O. loti*

The volatiles of adults and second instar nymphs were collected using solid-phase microextraction (SPME). For the initial use, the SPME fiber (57348-U, Supelco, Poole, UK) was inserted into the inlet of the GC at 250°C for a pretreatment duration of 30 minutes. Ten active second-instar nymphs or adult males or females were transferred to 1.5ml screw thread vials (J&K Scientific, Beijing, China) at room temperature, followed by exposing the SPME fiber to the headspace volatiles for 30 minutes. Headspace volatiles obtained in blank screw thread vials served as control. Three biological replicates were used for the experiment, and all tested thrips were used only once.

### 3.3 Analysis of headspace volatiles

The collected samples were analyzed using a Shimadzu GC-MS QP2010 PLUS with an HP-5MS column (30m × 0.25mm i.d. × 0.25μm film thickness; Agilent, Santa Clara, CA, USA). The carrier gas, helium, was set at a speed of 1.0 ml min^−1^. The injection temperature was 250°C. The oven temperature was held at 30°C for 1 min, then programmed to increase at 5°C min^−1^ to 150°C for 1 min, increase at 10°C min^−1^ to 200°C for 1 min, and finally increase at 20°C min^−1^ to 250°C for 1 min. Compounds were identified by comparing their mass spectra with NIST database. Moreover, synthetic standards were analyzed under the same conditions, supported by comparison of retention times and mass spectra.

### 3.4 Olfactometer assays

(R)-lavandulyl (R)-2-methylbutanoate and enantiomers were synthesized by the Laboratory of Organic Chemistry, College of Science, China Agricultural University, and (-)-Germacrene D was purchased from J&K Scientific, Beijing, China. The synthesized lavandulyl-2-methylbutanoate and the four enantiomers were first dissolved in paraffin oil and then diluted to a concentration of 0.5 μg/μl. Further, synthetic (R)-lavandulyl (R)-2-methylbutanoate was first dissolved in paraffin oil and subsequently diluted to concentrations of 0.5, 1, and 2 μg/μl. Synthetic (-)-Germacrene D was first dissolved in paraffin oil and subsequently diluted to concentrations of 1ng/μl, 10ng/μl, 50ng/μl, and 100ng/μl.

The responses of male and female *O. loti* to synthetic standards were tested in a Y-tube olfactometer. The glass Y-tube was made up of a stem and two arms (10 cm long and 1 cm in diameter) separated by a 75° angle from each other. Air flowing through the olfactometer was first purified with activated charcoal, then in the gas-washing bottle wetted with distilled water and split into two streams, each of which was fed through a 100 ml glass flask (test or control odor sources) and into one arm of the olfactometer at a flow rate of 50 ml min^−1^. Connections between the components of the olfactometer apparatus were made with Teflon tubes. Light for the olfactometer experiments was provided by a LED light (28W) placed at the Y-end of the olfactometer in a dark room at 26±1°C. Synthetic standards were dissolved in paraffin oil as the test odor source, with the control sample being paraffin oil. A single *O. loti* was introduced to the stem of Y-tube and observed for 5 min. When the test thrip entered a half-length of either arm of the Y-tube and stayed for 1 min, the choice was recorded. ‘No choice’ was recorded if a test thrip did not make enter an arm during the bioassay period. After five test thrips were tested, the location of the odor sources was swapped to avoid directional effects in the apparatus. And after 10 thrips tests, the Y-tube was cleaned with 75% ethanol and dried in an oven at 60°C for 30 minutes. Sixty biological replicates were carried out for each female or male treatment. Results were analyzed by Chi-square test using SPSS 22 (IBM, Armonk, NY, USA). ‘‘No choices’’ were excluded from the analysis.

### 3.5 Electroantennography assays

The insect antennal potential measurement system (IDAC2, Syntech, Kirchzarten, Germany) was applied to detect the electrophysiological response of adult male and female *O. loti* to synthetic (R)-lavandulyl (R)-2-methylbutanoate. The glass electrodes used were made of P-97 Flaming/Brown™ type micropipette puller (Sutter, Novato, CA, USA) and the electrolyte was 0.1 mol/L KCl solution. The antennae of active male and female adults of 3-5 days of age were dissected and the most proximal end was placed in the reference electrode and the two antennae in the recording electrode. Ten microliters of synthetic standard were added to the filter paper strip (0.5 × 5 cm) and inserted into glass Pasteur tubes for testing, and the control sample was liquid paraffin oil. Synthetic standard was first dissolved in paraffin oil and subsequently diluted to concentrations of 0.5, 1, and 2 μg/μl. Ten biological replicates were conducted in each treatment. The control data was obtained from the mean value of the two measurements at the beginning and the end. Results were recorded and processed by EAGPro 2 software. The student’s *t*-test was used to compare responses to odors and controls.

## Discussion

Aggregation is a common phenomenon in Thysanoptera and has been observed in different organs of host plants [22,28,29], demonstrating that aggregation pheromones may be present to induce aggregation behavior. To date, aggregation pheromones have been identified in six species of Thripidae, which could attract both male and female adults [19-21,23-25]. In our study, characterization of the headspace volatiles of the male *O. loti* revealed the presence of an ester named (R)-lavandulyl (R)-2-methylbutanoate. This is the first report of a male-produced aggregation pheromone in the genus *Odontothrips*. More notably, the aggregation pheromone (R)-lavandulyl 3-methylbutanoate, identified in *M. sjostedti*, is structurally similar to the substance in our study [23]. Phylogenetic analysis based on cladistics revealed that *Odontothrips* was most closely related to *Megalurothrips*, which partially explains the structural similarity of the aggregation pheromone of *O. loti* and *M. sjostedti* [30]. Further, phylogenetic analyses also explained the extent of variation in aggregation pheromone structure being found in other known thrips. Even two species of thrips in the same genus share the same aggregation pheromone components, but differ only in the proportions released, allowing to precisely differentiate between them [19,20].

Interestingly, we found that the male-specific aggregation pheromone of *O. loti* is likewise a minor sex pheromone for female adults of *Phenacoccus madeirensis* (Hemiptera: Coccoidea: Pseudococcidae), which is a polyphagous pest [31]. In addition, an enantiomer of this component has been reported to function as sex pheromones in Pseudococcidae [32]. This sharing of the same pheromone by two apparently distantly related species does not appear to be a coincidence, but reasonable evidence is missing. It may be relevant that host plant species of *P. madeirensis* [33] do not overlap with *O. loti*.

Mound (2009) [34] demonstrates that pore plates widely occur on the abdomens of Thripidae males and that their shape varies greatly between species. As well, we observed one 0.8-μm-diameter pore on each of the ventral sternites of the ?-? abdominal segments of adult males of *O. loti* (Figure S1), which is consistent with the location of the known reported pore plates. Intensive research on these pores has discovered that the glandular tissue is consistent with pheromone production, thus indirectly proving that pheromones are released by these areas [35,36]. If pore plates do produce aggregation pheromone, then aggregation pheromones are likely to be widespread, at least in the Thripidae. However, such a hypothesis has been complicated with the discovery of multiple pheromones. after all, it is not yet clear evidence that pore plates produce aggregation pheromones.

The olfactometer bioassays conducted in this study gave behavioral results show that both male and female *O. loti* adults were attracted to the synthetic (R)-lavandulyl (R)-2-methylbutanoate, indicated that the pheromone produced by *O. loti* males is an aggregation pheromone. That males are more sensitive to aggregation pheromones than females may be attributed to the greater activity of males [18], a phenomenon consistent with that reported for other thrips [24]. Previous studies have illustrated the efficacy of thrips aggregation pheromones combined with sticky traps in the field. Specifically, in a strawberry field, blue sticky traps baited with neryl (*S*)-2-methylbutanoate doubled trapping capacity and reduced adult thrips numbers per flower by 73% [37]. The same deployment in tea plantations resulted in significant reductions in adult *D. minowai* populations per 100 leaves, ranging from 29% to 59% [25]. Thus, the utilization of aggregation pheromones provides a basis for optimizing mass capture or integrating them with other control methods.

In our study, a larvae-specific compound called (-)-Germacrene D was identified, to which adult males and females did not respond significantly. Germacrene D is naturally occurring in plants, bacteria, and fungi [38-41] and may be active as an antibiotics, repellents, attractants, or pheromones, or may be produced to attack herbivores [40,42]. Nevertheless, the release of this sesquiterpene by insects has rarely been reported, and its role in populations of *O. loti* needs to be further investigated.

In conclusion, the male-specific aggregation pheromone (R)-lavandulyl (R)-2-methylbutanoate has been identified in *O. loti*, a component that significantly attracts both male and female adults. Further research will be devoted to field trials. As thrips are less able to fly upwind for long distances than other insects, such as moths, these percentage increases in response to pheromones are relatively low. Many attempts still need to be undertaken to improve trapping efficiency in the field, e.g. rational trap distribution, optimal trap dose, combined with other measures, including entomopathogenic fungi [15], natural enemies [43], etc.

