## Supplementary material for "Identification of Aggregation Pheromone in *Odontothrips loti* (Thysanoptera: Thripidae)": Supplementary Materials.docx

**A**

**B**

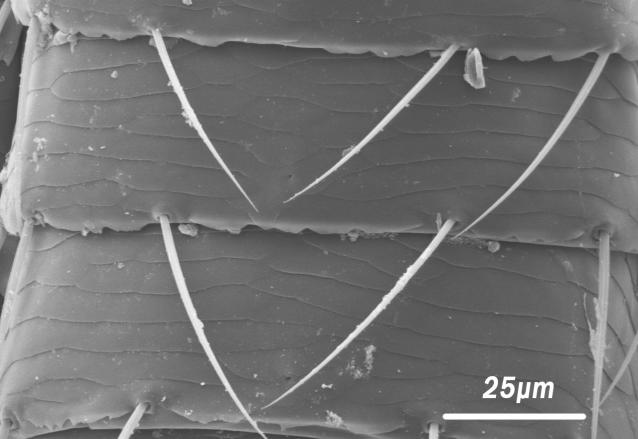

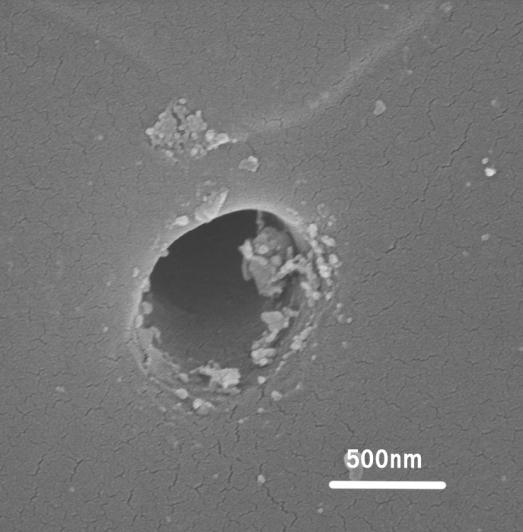

**Figure S1**. An overview of the pore plates on the adult males of *Odontothrips loti* under Scanning electron microscopy (SEM). (A) The pore plates (red arrow) are located at the Ⅵ-Ⅶ abdominal segments of adult males of *O. loti* and shown at higher magnification (B).
